# KRSA: Network-based Prediction of Differential Kinase Activity from Kinome Array Data

**DOI:** 10.1101/2020.08.26.268581

**Authors:** Erica A. K. DePasquale, Khaled Alganem, Eduard Bentea, Nawshaba Nawreen, Jennifer L. McGuire, Faris Naji, Riet Hilhorst, Jaroslaw Meller, Robert E. McCullumsmith

**Author notes:** co-first Authors (Erica A. K. DePasquale, Khaled Alganem).

## Abstract

**Motivation:** Phosphorylation by serine-threonine and tyrosine kinases is critical for determining protein function. Array-based approaches for measuring multiple kinases allow for the testing of differential phosphorylation between conditions for distinct sub-kinomes. While bioinformatics tools exist for processing and analyzing such kinome array data, current open-source tools lack the automated approach of upstream kinase prediction and network modeling. The presented tool, alongside other tools and methods designed for gene expression and protein-protein interaction network analyses, help the user better understand the complex regulation of gene and protein activities that forms biological systems and cellular signaling networks.

**Results:** We present the Kinome Random Sampling Analyzer (KRSA), a web-application for kinome array analysis. While the underlying algorithm has been experimentally validated in previous publications, we tested the full KRSA application on dorsolateral prefrontal cortex (DLPFC) in male (n=3) and female (n=3) subjects to identify differential phosphorylation and upstream kinase activity. Kinase activity differences between males and females were compared to a previously published kinome dataset (11 female and 7 male subjects) which showed similar patterns to the global phosphorylation signal. Additionally, kinase hits were compared to gene expression databases for *in silico* validation at the transcript level and showed differential gene expression of kinases.

**Availability and implementation:** KRSA as a web-based application can be found at http://bpg-n.utoledo.edu:3838/CDRL/KRSA/. The code and data are available at https://github.com/kalganem/KRSA.

**Supplementary information:** Supplementary data are available online.

## 1. Introduction

Protein phosphorylation marks one of the most important biological mechanisms that underlies various normal cellular functions, acting in complex protein-substrate networks. Phosphorylation cascades are also perturbed in many disease states (Hanahan and Weinberg, 2011; Simpson et al., 2019). As a result, kinases are one of the most studied proteins given their central role in normal and abnormal cell biological mechanisms (Ardito et al., 2017; Lahiry et al., 2010; Pawson and Scott, 2005; Ubersax and Ferrell, 2007). Kinomics, or the study of kinases and kinase signaling, has expanded from individual activity assays, with one peptide to study one kinase, to array or chip-based technology of up to 1000 reporter peptides, called kinase arrays or kinome arrays (Baharani et al., 2017; Diks et al., 2004; Houseman and Mrksich, 2002). The selected reporter peptides are designed to cover a broad range of signaling pathways, with large numbers allowing for a better understanding of kinase interactions and global changes that occur between two states (i.e., disease, cell type). However, analyzing the data from these peptide arrays is a complex process given that many kinases can phosphorylate the same peptide and an individual kinase can phosphorylate many peptides, making accurate interpretation of the data a challenging task. As the use of these kinome arrays becomes more widespread, there is an increasing need for tools that efficiently and accurately analyze these datasets. In particular, analytic tools are needed for nonexpert users of kinome array platforms.

Bioinformatics tools that are specifically designed to analyze kinome array datasets are beginning to emerge. One of these analytic tools is the Kinomics Toolkit, which gives users a platform for exploration of the peptide phosphorylation data, but does not provide upstream kinase predictions (Dussaq et al., 2018). This application pre-processes data, serves as a public data repository, and provides developers the opportunity to engage with the data in an SDK (software development kit). However, freely accessible tools for analyzing these kinome array data are relatively rare, with most research in this field relying on a mixture of manual statistical analysis, proprietary software such as BioNavigator by PamGene, or collaboration with bioinformatics experts. Another tool that was designed specifically to process kinase array data is the PamgeneAnalyzeR package, though this package is primarily focused on the pre-processing steps of kinase array datasets and not the downstream analysis (Bekkar et al., 2020).

Prediction of upstream kinase activity and network-based analyses provides a biologically-meaningful springboard for further research that is currently lacking in a user-friendly application. There are existing tools that aim to predict upstream kinases based on an input of enriched genes or phosphopeptides, like KEA (Lachmann and Ma’ayan, 2009) and PTM-SEA (Krug et al., 2019). However, none of these tools are specifically designed to take raw data from kinome array datasets and run a complete analysis pipeline starting from pre-processing to visualizing kinome networks.

A common and validated approach to predicting upstream kinase activity is to analyze the differences between 1) kinases predicted to be upstream of the peptides that are differently phosphorylated between two conditions and 2) kinases predicted to be upstream of the remaining peptides on the chip (Anderson et al., 2014; Isayeva et al., 2015). In a similar statistical approach, we have previously described a method which uses random sampling to identify highly active kinases from kinome array data (Bentea et al., 2019; Dorsett et al., 2017; McGuire et al., 2017). Briefly, we look at overrepresented/underrepresented kinases relative to an expected distribution using random permutation sampling of peptides. This type of analysis is valuable because it separates kinases that are truly differentially active from those who are highly active globally and don’t represent a change between states.

Here we present the Kinome Random Sampling Analyzer, or KRSA, which automates many of the steps described above, including peptide filtering, random sampling, heatmap generation, and kinase network generation. This new software makes analyzing kinome array datasets accessible and eliminates much of the human workload that the previous method required. More importantly, KRSA represents the results in a bigger biological context by visualizing altered kinome signaling networks instead of individual kinases. By designing this application with the web-based Shiny platform in R, we have made KRSA with a graphical user interface that is interactive and customizable, and open to timely updates as well as future expansion. KRSA can be used by biologists and data scientists alike, with no knowledge of statistical software required. This all-in-one tool is designed to move raw data from the kinome array through the processing pipeline to interaction network generation, creating downloadable figures and tables along the way to facilitate simple integration into publications and presentations.

This method has been applied to multiple datasets and predictions have been experimentally validated in our laboratory through individual kinase activity assays and inhibitor studies (Bentea et al., 2019; Bentea et al., 2020; Dorsett et al., 2017; Flaherty et al., 2019; McGuire et al., 2017; Schrode et al., 2019). An early version of KRSA, containing only the random sampling algorithm, identified altered phosphorylation of peptides and subsequently perturbed kinase activity in the anterior cingulate cortex (ACC) between schizophrenia and control subjects (McGuire et al., 2017). This tool was also used to analyze date from frontal cortex and hippocampus of rats subjected to lateral fluid percussion as a model of traumatic brain injury (TBI) and their sham surgery counterparts to identify differences in kinase activity in these brain regions (Dorsett et al., 2017). We used the platform to explore the kinase activity in cortical neurons differentiated from induced pluripotent stem cells (iPSCs) from a schizophrenia patient with a 4-bp mutation in the DISC1 gene (Bentea et al., 2019). KRSA also was used to analyze kinome signatures of genetic perturbation of NRXN1 and FURIN1 in human induced pluripotent stem cell (hiPSC)-derived neurons (Flaherty et al., 2019; Schrode et al., 2019). More recently, KRSA has been utilized to analyze the kinome signature of mice with a genetic deletion of a specific subunit of cystine/glutamate antiporter system (xCT −/− mice) (Bentea et al., 2020). All of these published studies required significant use of multiple tools, some closed-access, and statistical expertise for quality control, network growth, pathway enrichment, and manually created graphs, heatmaps, and networks for visualization purposes.

Interest within the neuroscience community in defining sex differences in the brain has increased over the past several decades. Differences in kinase activity and signaling between males and females have been implicated in sex-related variations in neuronal cell survival, outcomes after brain injury, and fear extinction, among other research areas (Armstead et al., 2017; Matsuda et al., 2015; Zhang et al., 2003). While these studies increased our confidence that we would be able to identify sex-based differences in kinase activity through the KRSA algorithm, we expected that these differences in kinase activity would be small compared to previous experiments we performed in the areas of TBI and schizophrenia. We also paired this experiment with a previously published kinome array study to compare against our findings (Rosenberger et al., 2016).

Indeed, we chose this research area to illustrate the use of KRSA and the importance of increasingly sophisticated bioinformatics tools in cases where the effect size is small and there are many confounding factors that limit interpretability. KRSA, in conjunction with other tools and methods designed for different steps in the gene expression process, can better elucidate the complex regulation of gene and protein activity that forms the basis of diversity.

## 2. Systems and Methods

### 2.1 Kinomics

#### 2.1.1 Sample preparation

For comparing kinase activity levels between female and male dorsolateral prefrontal cortex (DLPFC), we analyzed postmortem tissue obtained from 3 male and 3 female control subjects (for demographics, see Supplementary Table S1). Samples were lysed on ice for 30 minutes using M-PER lysis buffer (ThermoFisher) containing 1:100 Halt Protease and Phosphatase Inhibitor Cocktail (ThermoFisher), and centrifuged (14000 rpm, 10 min, 4oC). The supernatants were collected and assayed for total protein concentration (Pierce BCA Protein Assay Kit, ThermoFisher). Multiple aliquots of each sample were stored at -80oC. As freeze-thaw cycles can result in loss of kinase activity, frozen aliquots were used only once for kinase activity determination (Hilhorst et al., 2013).

#### 2.1.2 Serine-threonine kinase activity data generation

Profiling of serine-threonine kinase (STK) activity was performed as described in Bentea et al. (Bentea et al., 2019). Briefly, the activity assay was performed using the PamStation12 microarray (PamGene International) and STK PamChips containing 144 consensus phosphopeptide sequences (142 unique sequences, 2 internal controls) per array. Each array was blocked with 2% bovine serum albumin (BSA) before 2 µg of protein, 157 µM adenosine triphosphate (ATP), and a primary antibody mixture as a part of two-step reaction process. For the second step, FITC-labeled anti-phospho serine-threonine antibodies (PamGene) were added to each array. To facilitate interaction between kinases in the sample and specific peptides immobilized on the chip, the samples containing the active kinases and assay mix were pumped through the array. The phosphorylation levels, represented in this assay by the level of fluorescence in each array, were measured using Evolve kinetic image capture software (PamGene). Every 6 seconds for 1 hour the software captures FITC-labeled anti-phospho antibodies binding to each phosphorylated peptide. Peptide label intensity was captured at 10, 20, 50, 100, 200 ms during the post-wash phase of the procedure.

#### 2.1.3 Data analysis and filtering

Linear regression slope of the signal intensity as function of exposure time was used to represent the peptide phosphorylation intensity for downstream comparative analyses, averaged across the 3 biological replicates. This is done to increase the dynamic range of the measurements. The signal ratio between female and male DLPFC was used to calculate fold change (FC) values. Peptides with a fold change of at least 35% (i.e. FC > 1.35 or FC < 0.65) were considered differentially phosphorylated for the purposes of using KRSA. This threshold was chosen based on previous reports that suggest small changes in kinase activity are sufficient to trigger biologically relevant changes (Appuhamy et al., 2014; Dorsett et al., 2017; McGuire et al., 2017). Peptides that had very low signal or an R_2_ of less than 0.90 during the corresponding linear regression were considered undetectable or non-linear in the post-wash phase and were excluded from subsequent analyses.

#### 2.1.4 Heatmap and global phosphorylation plots

For generating the peptide phosphorylation heatmap and comparing the global phosphorylation levels, the linear regression slope of each peptide was multiplied by 100 and log_2_ transformed (Dorsett et al., 2017; McGuire et al., 2017). This interactive heatmap within the KRSA application is sorted in decreasing order of fold change between the given conditions.

#### 2.1.5 Independent kinome dataset

We analyzed a previously published kinome array dataset that studies the changes in protein kinase activity during Alzheimer’s Disease (AD) pathogenesis (Rosenberger et al., 2016). This postmortem study was performed using hippocampal (HPC) brain section samples. From this dataset, we reanalyzed all of data for the control samples (Braak Stages 0-1) for both female and male subjects. Given the Braak Stage 0 samples only contains male subjects and the apparent effect of Braak Staging on the kinome signatures, we limited ourselves to samples with Braak Stage 1, and that resulted into having 18 subjects (11 female and 7 male) to compare our results. We pre-processed this dataset identically to our established pre-processing methods, both for detectability and linearity of the slope in the post-wash phase. We generated a heatmap with unsupervised clustering and global phosphorylation plots. Given we have a higher number of samples in this dataset, we performed a principal component analysis (PCA) to cluster the dataset and also determine the factors that most explain the variance in these kinome signatures.

#### 2.1.6 Mapping upstream kinases

Protein kinases predicted to act on phosphorylation sites within the array peptide sequences were identified using GPS 3.0 and Kinexus Phosphonet (Kinexus Bioinformatics) (Xue et al., 2010). These programs provide predictions for serine-threonine kinases targeting peptide sequences ordered by likelihood of binding. The union of the highest ranked 5 kinases in Kinexus and kinases with scores more than two times the prediction threshold in GPS 3.0 were considered predicted kinases for each peptide and used in KRSA analysis (Bentea et al., 2019). This list was combined with kinases shown in the literature to act on the phosphorylation sites of the peptides via PhosphoELM (http://phospho.elm.eu.org) and PhosphoSite Plus (https://www.phosphosite.org). While the reporter peptides list remains largely stable across different versions of kinase array chips, some peptides are removed and replaced by the manufacturer to improve kinase predictions. Updates to the kinase-peptide list should be made to reflect the assortment of peptides on the new chip prior to running KRSA in accordance with the above guidelines. The user also has the option to upload their own protein-substrate mapping files based on phosphosite found in the chip to perform the upstream kinase analysis.

#### 2.1.7 Waterfall plots and individual peptide phosphorylation curves

In addition to the graphs and heatmaps automatically generated by the KRSA software, we used waterfall plots to illustrate the differences in phosphorylation across all peptides of the array in which the reporter peptides are displayed along the y-axis based on fold change values. Post-wash phosphorylation curves for individual reporter peptides were additionally plotted using the peptide spot intensities captured at 10, 20, 50, 100, 200 ms exposure times and averaged across the 3 biological replicates.

#### 2.1.8 Kinase network pathway analysis

We used pathway enrichment analysis with Enrichr (https://amp.pharm.mssm.edu/Enrichr/) to gain insight into biological pathways of the most prominent hits in using the kinase-kinase network produced by KRSA (Supplementary Table S2) (Chen et al., 2013). Results are based on the KEGG 2016 cell signaling pathway database (http://www.kegg.jp/) and ordered by the number of genes enriched in the pathway and significance of association.

### 2.2. Transcriptional validation

In order to identify whether the changes in kinase activity are mirrored by transcriptional changes in their corresponding genes, we looked for the kinase network genes (Supplementary Table S3) in two previously published region-level microarray databases of gene expression changes in female vs. male frontal cortex (Trabzuni et al., 2013; Xu et al., 2014). In addition, we probed the expression of the kinase network genes in a recently generated microarray dataset of laser capture micro-dissected (LCM) pyramidal neurons from deep and superficial DLPFC layers of control male and female subjects (Wu et al., 2020).

## 3. Algorithm

To determine which upstream kinases are most likely to be important in driving the changes in phosphorylation between female and male DLPFC, we performed random sampling analysis via KRSA. The major steps when running the application are as follows:

### 3.1 Inputs and Options

1. <u>Data Selection</u>: The user-supplied kinase-peptide association file and the raw kinome array data file are selected as input. The kinase-peptide associations should be based on the predicted interactions in GPS 3.0 and Kinexus Phosphonet, as described in ‘2.1.5 Mapping upstream kinases’. Group classifications for the input samples are determined through drop-down menus which allow for clustering of samples within and across chips via manually concatenated input files. Expected inputs should be formatted as shown in the example files on the project GitHub (https://github.com/kalganem/KRSA) and conform to standards imposed by leading laboratories generating PamChip kinase data. Descriptions of pre-processing requirements, software use, and downstream validation techniques can also be found at the associated project GitHub.
2. <u>Peptide Filtering Options</u>: Stringency options, including minimum exposure intensity > 2 at the last exposure time in the cycle (200 ms) and linearity of the post-wash curve as determined by linear regression (R_2_ > 0.9), are provided to reduce the total number of peptides evaluated to those meeting the quality control standards desired by the user, with these peptides denoted as *s*. Fold change thresholds are selected to identify differentially phosphorylated peptides (*h*, or “hits”) between selected groups. KRSA has a default cutoff of ± 35% as a fold change threshold, though modification should be made in cases where the bar for biological significance is substantially higher or lower. Line plots displaying the phosphorylation intensity in the post-wash phase for each sample are generated to allow for visual inspection of the selected peptides before proceeding to the next step to ensure linearity and sufficient magnitude of phosphorylation for each differentially phosphorylated peptide.

### 3.2 Empirical measures of statistical significance

3. <u>Random Sampling</u>: KRSA performs random sampling of the available peptides on the kinase array to get distributions for associated kinase-peptide interactions. The number of iterations of sampling (*i*), between 0-5000, is selected by the user, where higher *i* values increases the stability of the kinase predictions while also increasing the run time of the software. For each iteration *i*, the same number of peptides as *h* are randomly selected without replacement from *s* available peptides (these randomly selected peptides are denoted as *h*’).
4. <u>Kinase Mapping</u>: Predicted kinases are mapped to *h*’ using the default kinase-peptide associations file (or supplied by the user) and the number of appearances for each kinase is calculated. When the sampling is complete, a distribution for each kinase is determined from the mean number of times each kinase was predicted based on the randomly sampled peptides and the corresponding confidence interval. The number of times a kinase is mapped to *h* (the differentially phosphorylated peptides) is also determined, and comparison of this kinase count to the mean kinase count from sampling allows us to determine differential kinase activity through a Z-score > 2 (alpha-level = 0.025).

### 3.3 Outputs

5. <u>Tabular Results</u>: Results of the above calculations are provided in tabular form, sorted in decreasing order by Z-score. Kinases appearing at the top of this table are those that are most likely to have significant alterations in activity between the control and the experimental groups. This table can be directly saved within the KRSA software for inclusion in publications.
6. <u>Visualization of Distributions</u>: Histograms displaying the distribution of mappings for each kinase after random sampling are provided, with the *h* mapping (vertical line), *h*’ averages (bars), and *h*’ confidence interval (gray translucent region) overlaid. Histograms of kinases with *h* averages outside of the confidence interval in either direction are considered significantly altered and match the kinases shown in the tabular output.
7. <u>Heatmap of Differential Phosphorylation</u>: A user-customizable heatmap, generated using the R package ‘gplots2’, visualizing the fold-change differences for each peptide is provided and sorted in descending order. This heatmap can be saved as a PDF or image file directly within the KRSA software.
8. <u>Kinase Network</u>: A network diagram connecting *h* kinases to predicted interacting proteins or kinases is generated using the Search Tool for Retrieval of Interacting Genes/Proteins (STRING) database data and graphed with the ‘igraph’ package in R (Szklarczyk et al., 2017). The generated network represents the direct interactions between protein kinases identified from KRSA, as well as additional kinases indirectly connected to the original seed kinases. STRING was used for growing and connecting the kinase network by selectively adding interacting kinases with the highest confidence interaction score. Because of the highly interactive and repetitive structure of kinase activity, we weighted the nodes of the network based on the number of interactions found for each kinase in the network. Confidence thresholds and connection type options are provided to allow for more control over the resultant graph.

## 4. Implementation

### 4.1 KRSA Shiny Application

We developed the Kinome Random Sampling Analyzer (KRSA) as an R Shiny application to facilitate processing and interpretation of PamChip kinase array data. KRSA performs quality control filtering of peptides, random sampling to identify highly enriched kinases, and network expansion to grow the list of potentially affected kinases for pathway analysis (Fig. 1). The KRSA app includes sortable tables, interactive heatmaps, networks and graphs, as well as simple download options throughout for saving both processed data files and publication-ready images. The Kinome Random Sampling Analyzer is freely available at http://bpg-n.utoledo.edu:3838/CDRL/KRSA/, and all source code, documentation, and sample files are located at https://github.com/kalganem/KRSA.

**Figure 1.**
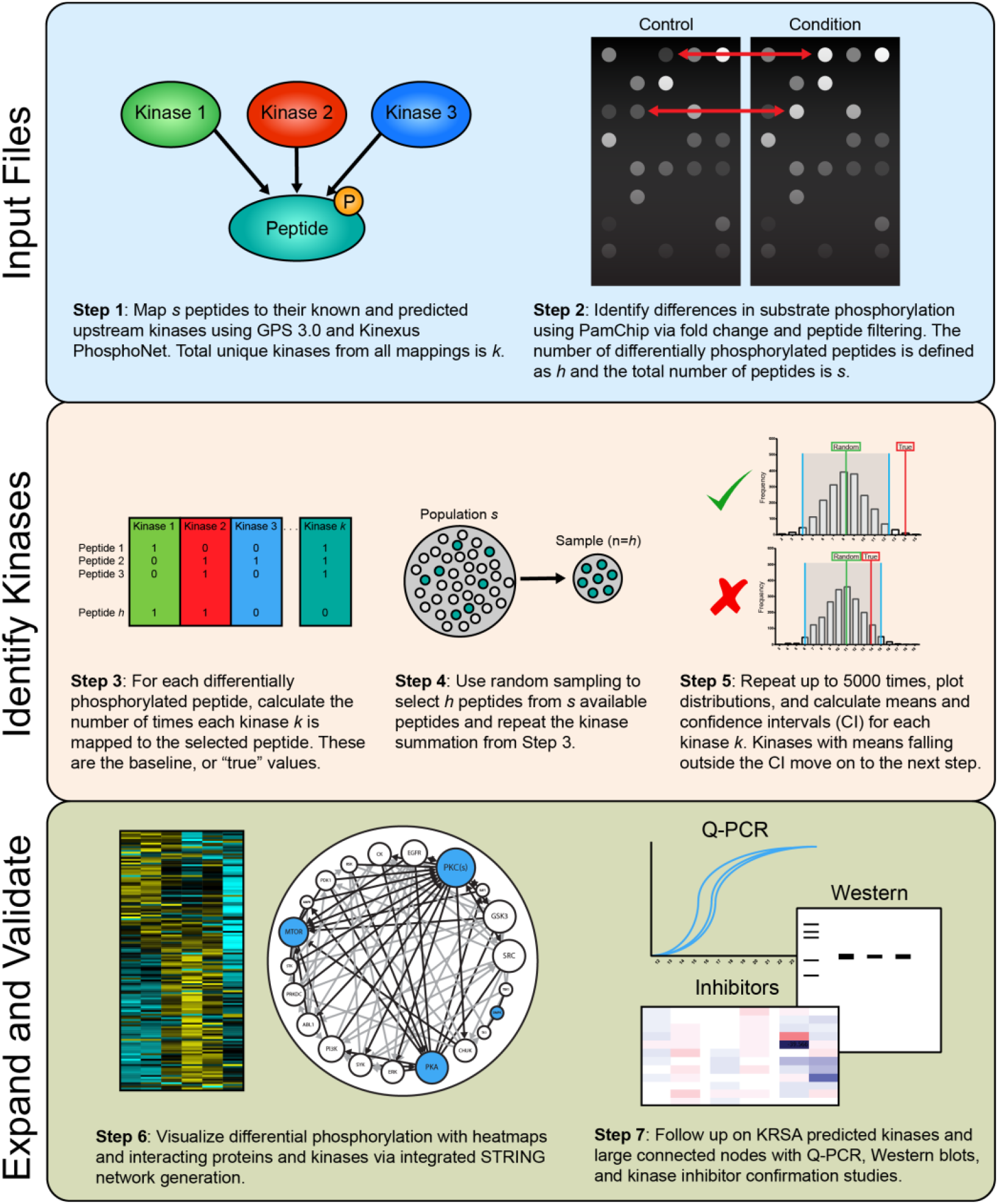
Workflow overview illustrating the primary steps of the KRSA pipeline. The “Input Files” section outlines the initial input files for KRSA, including the raw kinome array data and the peptide-kinase association file as well as the initial filtering step in KRSA. The “Identify Kinases” section describes the random sampling and distribution evaluation methods used to identify differentially active kinases. Finally, the “Expand and Validate” portion of the figure shows the kinase network generation step of KRSA and confirmation experiments that can be used to validate the predictions from KRSA.

### 4.2 Global serine-threonine protein kinase activity in female vs. male DLPFC

To elucidate differences in kinase activity between the brains of both males and females, we used KRSA to predict differential upstream kinase activity in conjunction with Enrichr for pathway analysis. KRSA filtered out 58 of the 144 peptides on the PamChip kinome array that were considered undetectable or were not linearly increasing with exposure time (based on R_2_ > 0.9). The signal at the remaining 86 reported peptides is depicted in a heatmap with phosphorylation intensity for each reporter peptide matched between the two sample groups Fig. 2A. The global phosphorylation levels (calculated as the average phosphorylation intensity across all reporter peptides) were not significantly different in female vs. male DLPFC (Fig. 2A inset; p = 0.8234 using a Mann-Whitney test).

**Figure 2.**
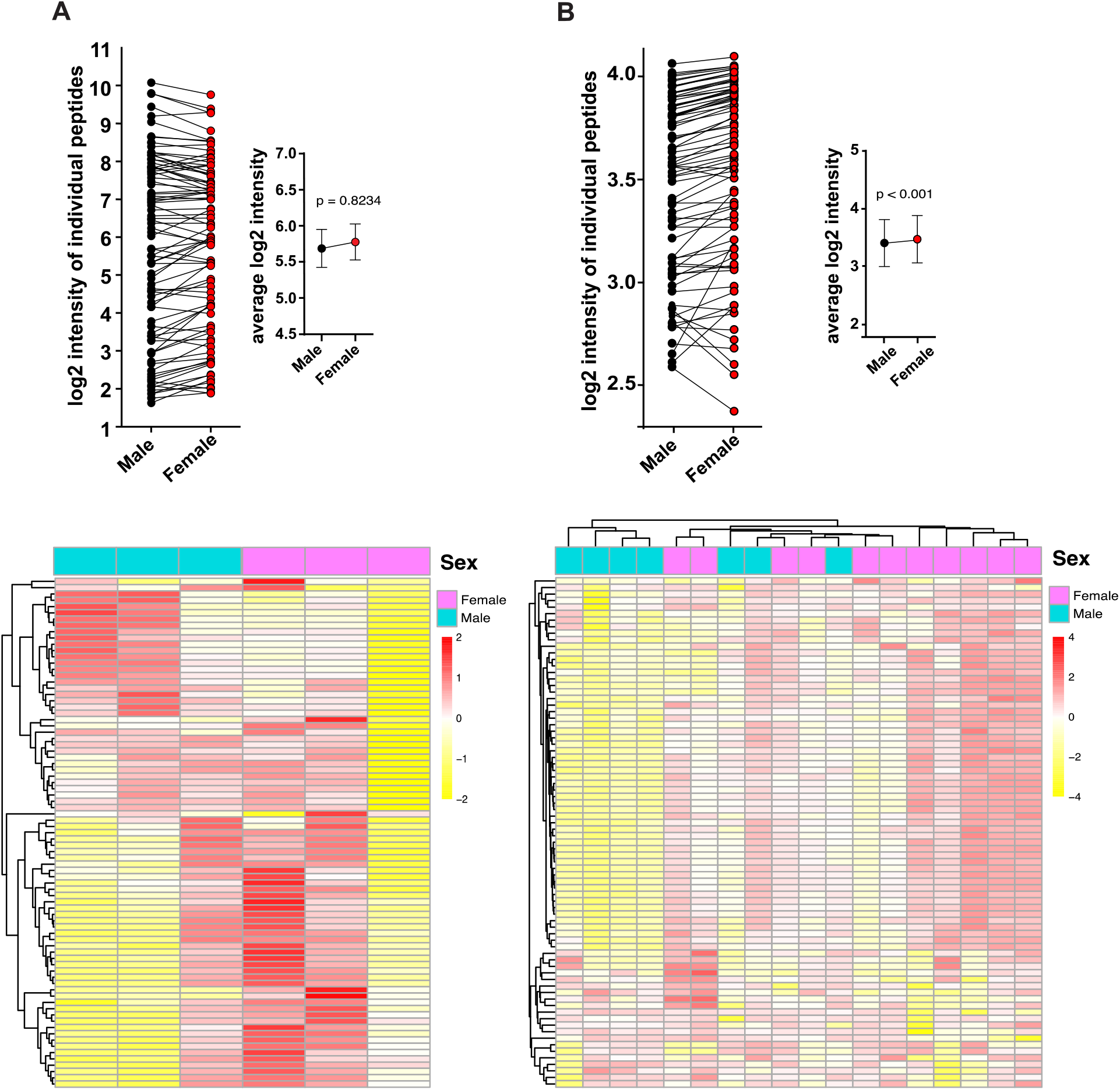
Global serine-threonine kinase activity in female vs. male DLPFC. (A) Global phosphorylation plots, showing changes in phosphorylation levels at each reporter peptide, as well as the average phosphorylation values (inset), when comparing female to male DLPFC. Data are presented as mean ± SEM and analyzed statistically using a Mann-Whitney test. Global phosphorylation heatmap generated by KRSA depicting the relative signal intensity at each reporter peptide for the 6 samples on the array (3 female and 3 male) (B) Global phosphorylation plots, showing changes in phosphorylation levels at each reporter peptide, as well as the average phosphorylation values (inset), when comparing female to male subjects using 18 control samples from the AD cohort (11 female and 7 male). Data are presented as mean ± SEM and analyzed statistically using a Mann-Whitney test. To highlight differences, the heatmap is normalized per row to present relative changes at each individual peptide between the groups. Red indicates relatively higher levels of phosphorylation and yellow indicates relatively lower levels of phosphorylation.

### 4.3 Global serine-threonine protein kinase activity of the independent dataset

To compare our observations against an independent kinome array study, we used the data of the control samples from the AD cohort (Rosenberger et al., 2016). In this dataset, all subjects were run using 5 to 6 technical replicates. For the analysis, all technical replicates were averaged (for demographics, see Supplementary Table S2). Similar to our own dataset, we generated both a heatmap representing global phosphorylation levels as well as averaged global signal for females and males. We observed a significant difference between global female vs. male HPC kinome signatures. (Fig. 2B inset; p < 0.001, using a Mann-Whitney test).

### 4.4 Altered kinase activity in female vs. male DLPFC

To examine only peptides with robust changes in phosphorylation, we restricted our selection of peptides to those with a fold change difference of ±35% between the sexes, which resulted in 22 peptides changed in phosphorylation between female vs. male DLPFC (Fig. 3A, red bars). Representative phosphorylation intensity graphs automatically generated by KRSA are given in Fig. 3B, with each line representing a biological sample and the colors indicative of the groups (male or female). Using the 86 peptides that remained following filtering, we performed random sampling analysis via KRSA to identify the upstream kinases predicted as differentially active and likely drivers of the differences observed in male vs. female brain (Dorsett et al., 2017; McGuire et al., 2017). This led to the identification of 7 different serine-threonine kinase families differentially represented in DLPFC (Table 1). Histograms are generated by KRSA for every potentially implicated kinase; Fig. 4 shows examples reflecting the results of the random sampling for kinases overrepresented in female vs. male DLPFC (CDK, PDK1, and P38, Fig. 4A-C; identified more than by random chance), as well as for kinases which were not differentially represented between the sexes (MAPKAP, MTOR, and GSK, Fig. 4D-F; identified as expected by random chance).

**Table 1.** Predicted kinases and distributions for female vs. male DLPFC.

| Kinase | Observed Hits | Distribution Mean | Standard Deviation | Z-score | Confidence Interval |
| --- | --- | --- | --- | --- | --- |
| CDK | 9 | 15.59 | 1.98 | 3.32 | 11.62 to 19.56 |
| PDK1 | 4 | 9.81 | 2.16 | 2.69 | 5.49 to 14.13 |
| P38 | 3 | 8.09 | 2.09 | 2.44 | 3.92 to 12.27 |
| BARK2 | 2 | 0.48 | 0.63 | 2.40 | -0.78 to 1.75 |
| BUD32 | 1 | 0.15 | 0.36 | 2.34 | -0.57 to 0.88 |
| WNK | 1 | 0.16 | 0.36 | 2.32 | -0.57 to 0.88 |
| MLK | 6 | 2.79 | 1.42 | 2.26 | -0.06 to 5.63 |

**Figure 3.**
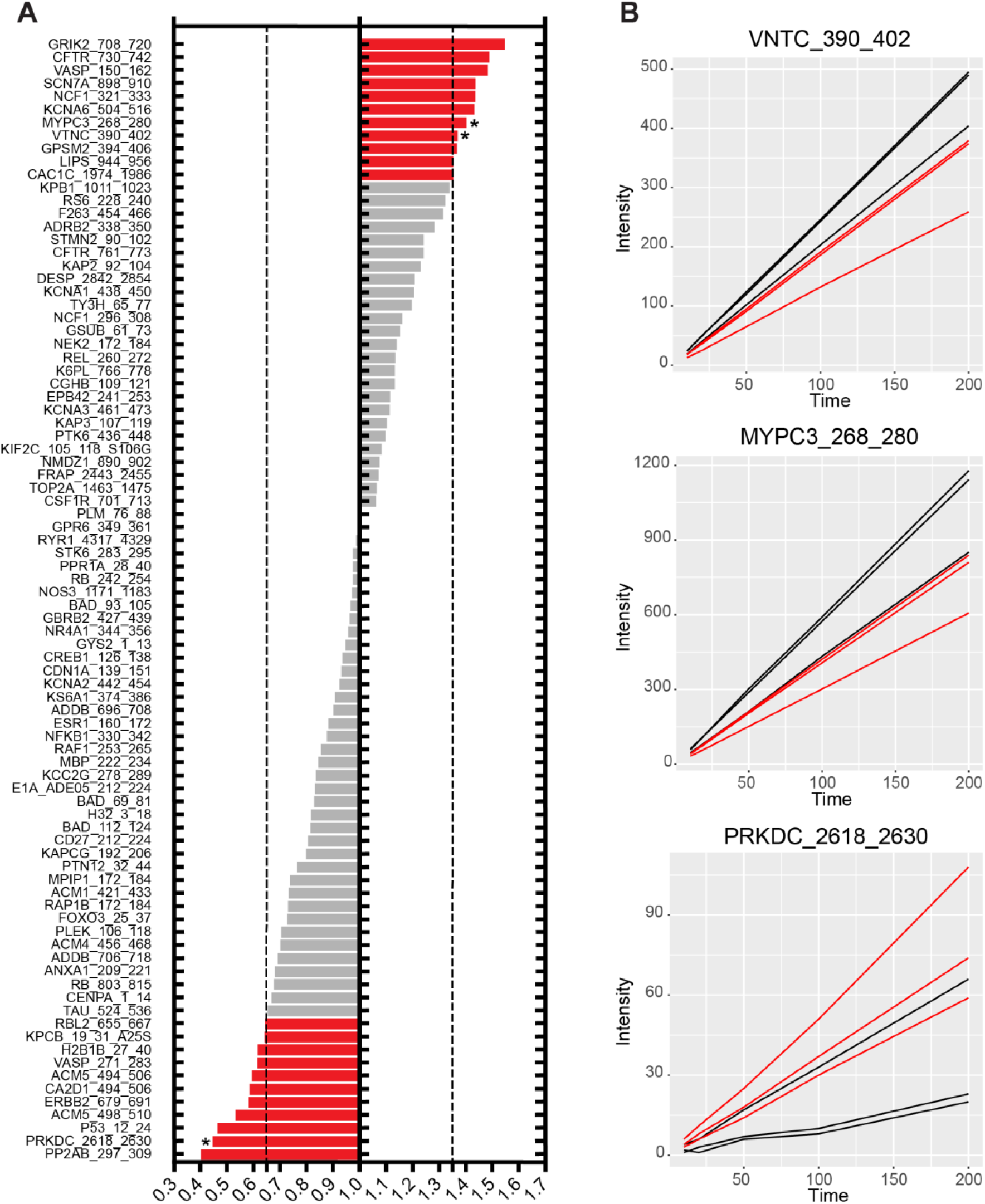
Changes in phosphorylation at reporter peptides in female vs. male DLPFC. (A) Waterfall plot showing changes in phosphorylation at reporter peptides for female vs. male DLPFC. Peptides with increased (FC > 1.35) or decreased phosphorylation (FC < 0.65) are highlighted in red at the top and bottom, respectively. (B) Representative examples of post-wash phosphorylation curves of reporter peptides in female vs. male DLPFC (marked with * in panel A).

**Figure 4.**
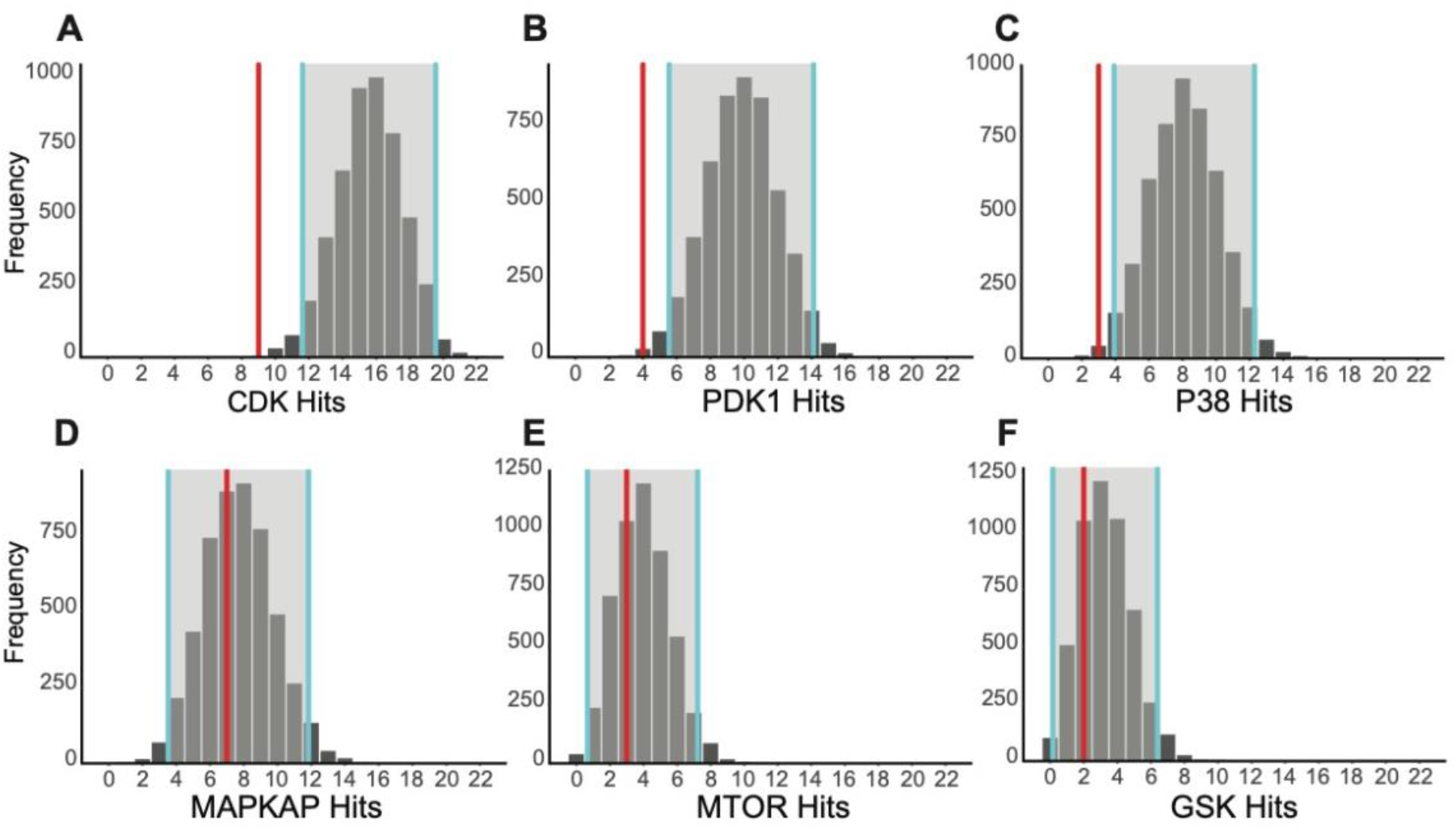
Observed frequency of selected kinases relative to expected random sampling distribution in female vs. male DLPFC. Examples are shown for kinases identified in the reporter peptides more than by random chance alone (CDK, PDK1, and P38; A-C), as well as for kinases identified as expected by random chance alone (MAPKAP, MTOR, and GSK; D-F). KRSA was performed with 5000 iterations and histograms were automatically generated. Gray areas between 2 blue lines indicate ± 2 standard deviations from the expected distribution mean. The prevalence of the selected kinase within the identified differentially phosphorylated peptides is indicated in red.

### 4.5 Altered kinase activity in female vs. male in the independent dataset

We used a similar method to determine the upstream kinase hits for the HPC cohort, and that analysis led to the identification of 5 different serine-threonine kinase families differentially represented in HPC between female and male control subjects (|Z score| >= 2). The list of kinase hits comprised of PKN, MLCK, SGK, TAO, and STE7 (Table 2).

**Table 2.** Predicted kinases and distributions for female vs. male Hippocampus (AD cohort, only control subjects)

| Kinase | Observed Hits | Distribution Mean | Standard Deviation | Z-score | Confidence Interval |
| --- | --- | --- | --- | --- | --- |
| PKN | 2 | 0.302 | 0.491 | 3.462 | -0.68 to 1.28 |
| MLCK | 3 | 0.768 | 0.802 | 2.782 | -0.84 to 2.37 |
| SGK | 8 | 3.888 | 1.694 | 2.428 | 0.50 to 7.28 |
| TAO | 1 | 0.146 | 0.353 | 2.423 | -0.56 to 0.85 |
| STE7 | 3 | 7.615 | 2.048 | 2.254 | 3.52 to 11.71 |
| AKT | 9 | 6.034 | 1.884 | 1.574 | 2.27 to 9.80 |
| PKCD | 0 | 1.937 | 1.239 | 1.563 | -0.54 to 4.41 |

### 4.6 Kinase network model of female vs. male DLPFC

The complexity of cellular signaling ensures that kinases do not act in isolation, but instead as part of an interacting network with other kinases and proteins that regulate biological processes (Yao et al., 2015). The nature of this system means that final KRSA predictions should include potentially interacting kinase families for downstream pathway analysis and hypothesis generation. To accomplish this goal, KRSA connects the initial set of 7 predicted kinases (Table 1) with kinase families that are known to interact with using the STRING database. Figure 5 depicts the extended kinase network including associated or indirect kinase interactions. This analysis revealed members of the CDK, PDK1, BARK2, P38, BUD32, WNK, and MLK kinase families as particularly large nodes of regulation of the female vs. male kinase network model. Enrichr was performed using the 103 genes that form the extended kinase network as input (any kinase with a Z score above 2) (Supplementary Table S3), which resulted in “MAPK signaling pathway”, “Neurotrophin signaling pathway”, “T cell receptor signaling pathway”, “FoxO signaling pathway”, and “TNF signaling pathway” as top cellular pathways associated with the female vs. male differential kinase network (Table 3).

**Table 3.**
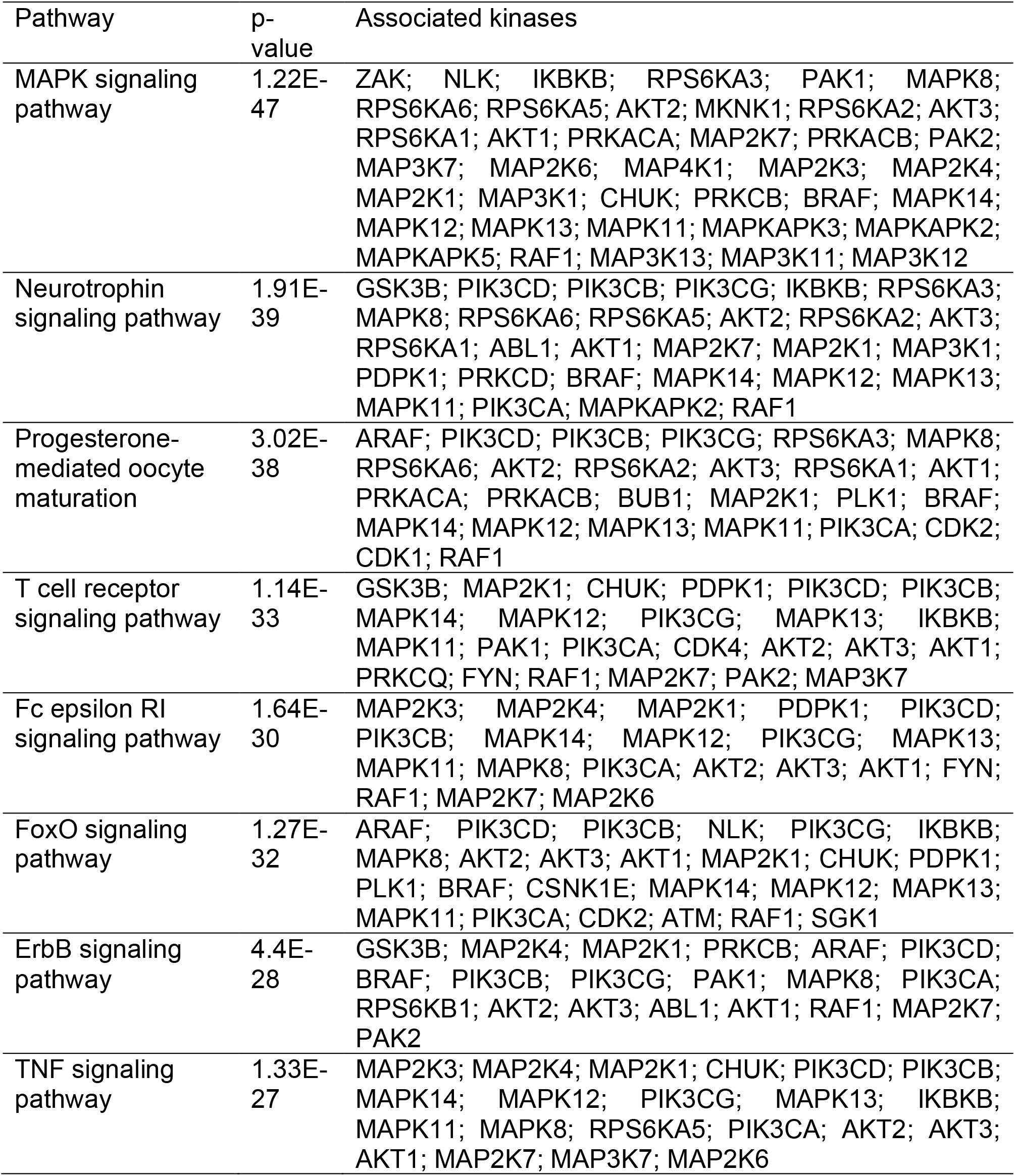
Enrichr cell pathway analysis (KEGG 2016) of the female vs. male kinase network.

**Figure 5.**
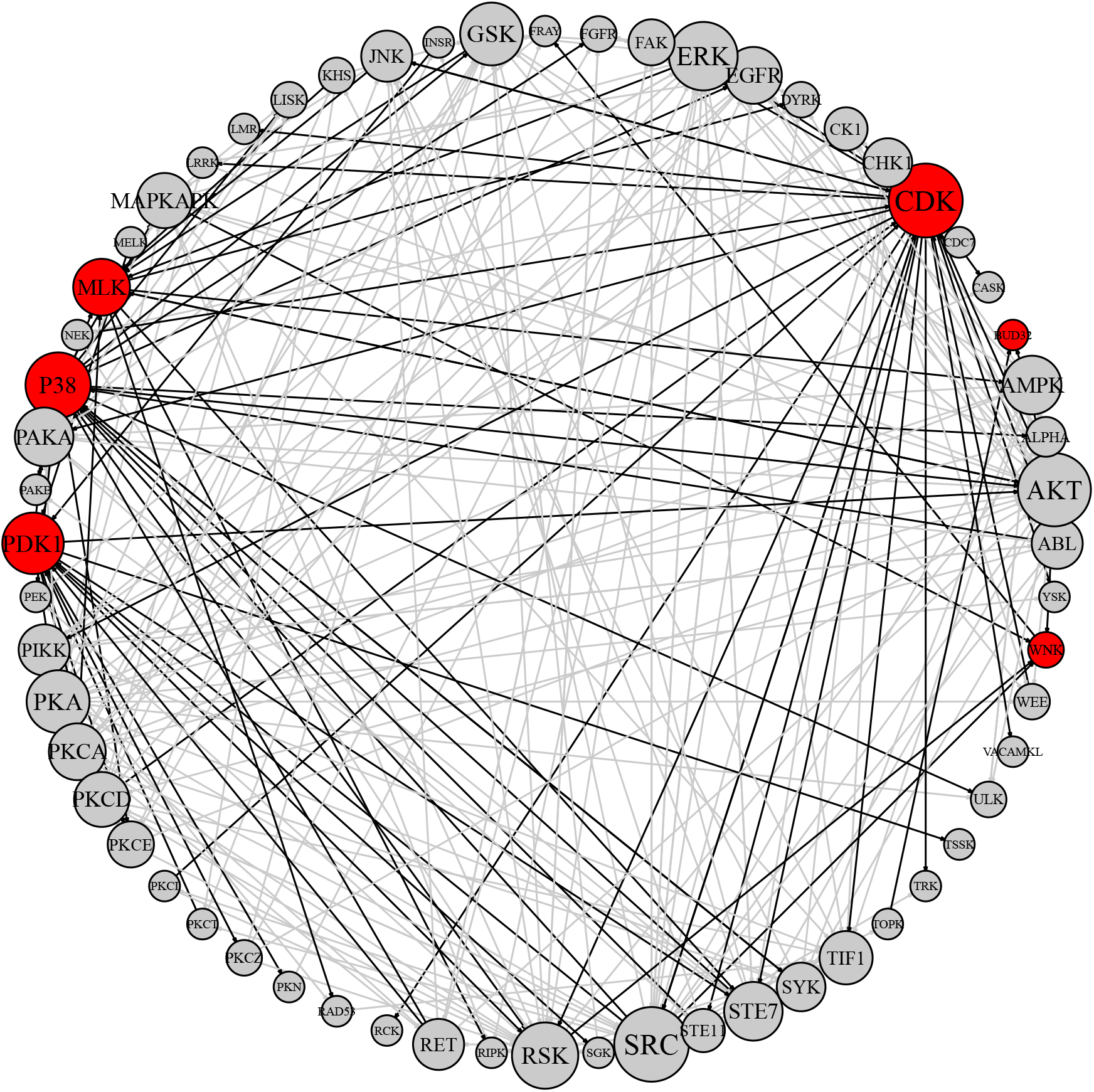
Kinase network model of female vs. male DLPFC. The kinase network was obtained in KRSA by growing the kinome array hits with kinase interacting partners as identified using STRING. The kinome array hits are color coded: red circles reflect kinase families identified in the kinome array and gray circles represent indirect or associated hits obtained after growing the network in STRING. Circle size corresponds to the number of interactions, with larger circles having more interactions. Black lines represent interactions with a kinome array direct hit, while gray represent interactions made between associated STRING-expanded kinase families.

### 4.7 Changes in kinase gene expression in female vs. male DLPFC

To identify whether the changes in kinase activity are mirrored by transcriptional changes in the corresponding genes, we queried the kinase network genes (Supplementary Table S3) in 2 previously published gene expression microarray databases of gene expression changes in female vs. male frontal cortex (region-level), as well as in a recently generated microarray dataset of LCM pyramidal neurons from deep or superficial layers of female vs. male DLPFC (unpublished data) (Trabzuni et al., 2013; Xu et al., 2014). This revealed a complex transcriptional signature with a large subset of genes showing changes in expression in female vs. male DLPFC pyramidal neurons, some of which extended at region-level (Supplementary Tables S4 and S5). These findings support the kinase dysregulation profile we identified using KRSA and suggest that some of the changes in kinase activity may be attributed to changes in kinase gene expression, though other factors play a large role like a global kinase activity.

## 5. Discussion

Alterations in kinase activity have been implicated in many diseases, both in and out of the nervous system, including cancers and neurological disorders (Dermit et al., 2017; McGuire et al., 2014; Zheng et al., 2012). The core KRSA algorithm, prior to transformation into a publicly available web-application, has been used to answer many research questions and generate new hypothesis by our research team and collaborators (Bentea et al., 2019; Dorsett et al., 2017; Flaherty et al., 2019; McGuire et al., 2017; Schrode et al., 2019). Recently, we identified altered AKT and JNK activity in schizophrenia by performing the core KRSA algorithm on postmortem anterior cingulate cortex (McGuire et al., 2017). Additional kinome arrays with targeted inhibitors, protein and phosphoprotein measurement, and specific kinase assays confirmed our predictions and supported the previously unreported finding of JNK activity alterations in schizophrenia (Morris and Pratt, 2014). The KRSA methodology was also applied to a rat model of traumatic brain injury which showed global glutamate transporter dysregulation in the brain mediated by region-specific differences in kinase activity (Dorsett et al., 2017). These predictions were validated through protein and phosphoprotein measurement, targeted inhibitors, and glutamate uptake assays. With the method experimentally validated, the goal was to design KRSA as an accessible web-based application that automates the entire process from raw data filtering to kinase network generation.

Unlike many diseases and conditions, where distinct high-magnitude changes in gene expression and subsequent downstream function occur because of the disease processes, differences between healthy male and female brains are theoretically subtler and harder to characterize. As an example of KRSA’s capabilities, we probed for kinase activity differences in the male and female brain using postmortem dorsolateral prefrontal cortex. KRSA predicted seven specific kinases to be different between males and females, which was grown to nine kinase families using the built-in STRING database. The kinase families and their members predicted by KRSA are key components of many signaling pathways, including MAPK, neurotrophin, T-cell receptor, FoxO, and TNF signaling, indicating that a large network of subtle differences exist between males and females.

We also compared our findings to a previously published kinome array study. The samples from that study showed a similar pattern of changes between female and male kinome signatures as an overall higher phosphorylation levels in the female samples. For the HPC cohort, the unsupervised clustering also showed a clear separation between male and female signatures. Given the higher number of samples in the independent dataset, we were able to statistically show a significant difference of kinome signatures between the two sexes (p< 0.001). Using Z-score threshold of 1.5, we saw an overlap of the STE7 family (MAPKK, MAP2K, MEK) among the two cohorts. Moreover, using Z score threshold of 1.25, we observe a bigger overlap with 6 kinase families (STE7, STE11, AMPK, HAL, PKN, PAKB) (Supplementary Figures S2).

These results highlight the need of separating male and female sample groups given the apparent difference of their kinome signatures. Experiments that combine male and female samples with other independent variables may be difficult to interpret, as some of the changes in the kinome may be due to sex differences and not, for example, disease state. This supplements the findings at the gene transcript level where many studies showed sex-specific differences in gene expression and gene regulatory networks (Lopes-Ramos et al., 2020; Trabzuni et al., 2013).

There are some limitations that may restrict the use of this application while at the same time leaving room for continued expansion and improvement. The magnitude and complexity of kinome array data, along with its relative uniqueness, has given rise to multiple methods for calculating intensity for the selection of significantly altered peptides, namely using end-level readings alone or in combination with kinetic data from the PamChip. While many laboratories have successfully used end-level readings to identify biologically relevant targets, this being the method that KRSA currently uses, Dussaq et al. notes that the kinetic data is under-utilized and has the potential to uncover additional relevant information. This is mainly applicable to the tyrosine kinase chip, but not the serine-threonine kinase chip used for this study (Dussaq et al., 2018). However, inclusion of these kinetic data could be implemented in KRSA in the next development version to expand the quality control and filtering capabilities. Another possible limitation is the relatively rigid requirements for the input data format, which may necessitate pre-processing of the raw datasets in order to fit the required format for the KRSA. To alleviate some of this concern, we have provided example format files on our GitHub page to enable users to convert their files into a KRSA-friendly format.

As research moves away from examining incremental biological steps in isolation and toward functional assays and whole systems-based approaches, experimental techniques like kinome chips and arrays will become more relevant and widely used. Bioinformatics has breached these areas, with cutting-edge tools being created for use primarily by other bioinformaticians and statisticians. In the area of kinomics, there is a need for end-to-end processing of kinome array data in a user-friendly, open source, and interactive environment. The Kinome Random Sampling Analyzer (KRSA) fills this gap in the field and will serve as a stepping stone for the use and interpretation of kinome array data for laboratory biologists and computational biologists alike.

## Acknowledgments

We would like to thank Daniel Schnell for helpful discussions on the statistical methods behind KRSA. We also thank the developers of R studio, Shiny, and supporting R packages used in the implementation of KRSA.

## Supplementary Data

**Supplementary Figure 1.**
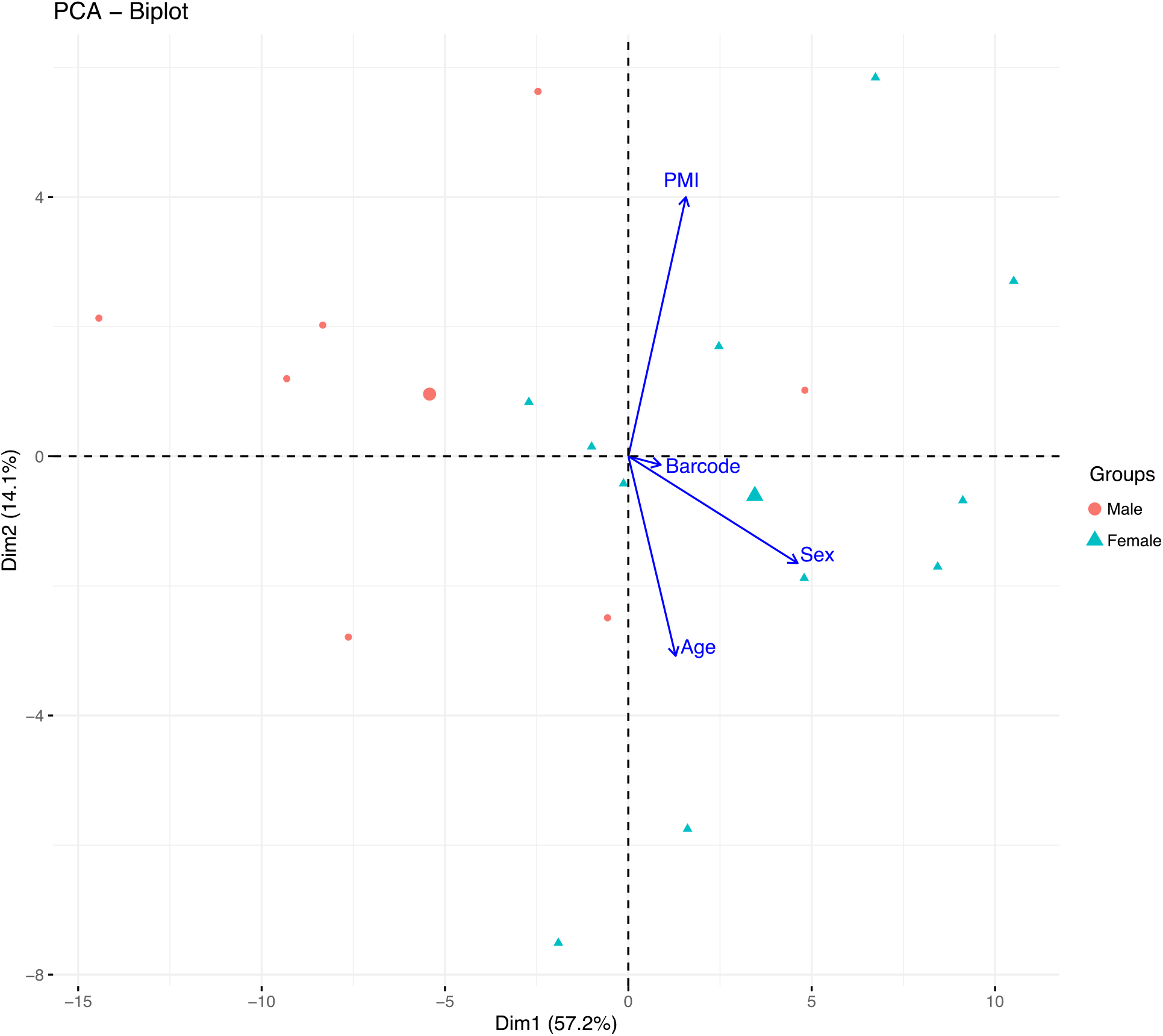
Principal Component Analysis (PCA) of the independent dataset from the AD cohort. Using the subjects in AD cohort dataset (controls only) showing the clustering of samples and the factors that most explain the variance in the kinome signatures. PMI: postmortem interval, Barcode: Chip ID.

**Supplementary Figure 2.**
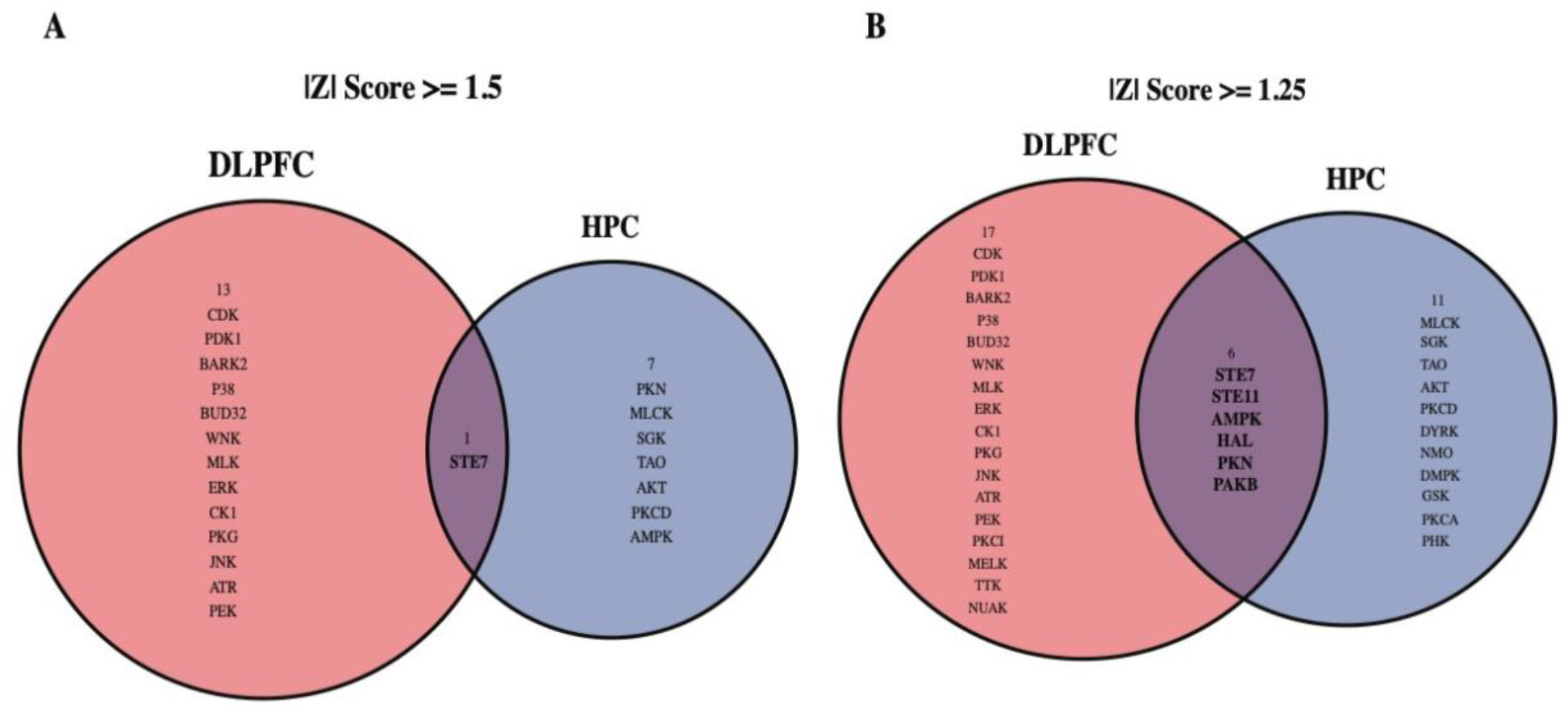
Venn diagrams showing overlap of the total of overrepresented/underrepresented kinases for both cohorts. DLPFC from current study, HPC from the AD cohort study (only control subjects). (A) Filtered kinase with absolute values of Z scores equal or above 1.5 for both datasets. (B) Filtered kinase with absolute values of Z scores equal or above 1.25 for both datasets. DLPFC: dorsolateral prefrontal cortex, HPC: hippocampus.

**Supplementary Table S1.** Kinome array subject demographics.

| Subject | Sex | Age | pH | PMI (h) |
| --- | --- | --- | --- | --- |
| M1 | M | 73 | 6.4 | 17 |
| M2 | M | 71 | 6.4 | 13 |
| M3 | M | 71 | 6.4 | 20 |
| F1 | F | 76 | 6.3 | 23 |
| F2 | F | 73 | 5.9 | 25 |
| F3 | F | 77 | 6.6 | 30 |

**Supplementary Table S2.**
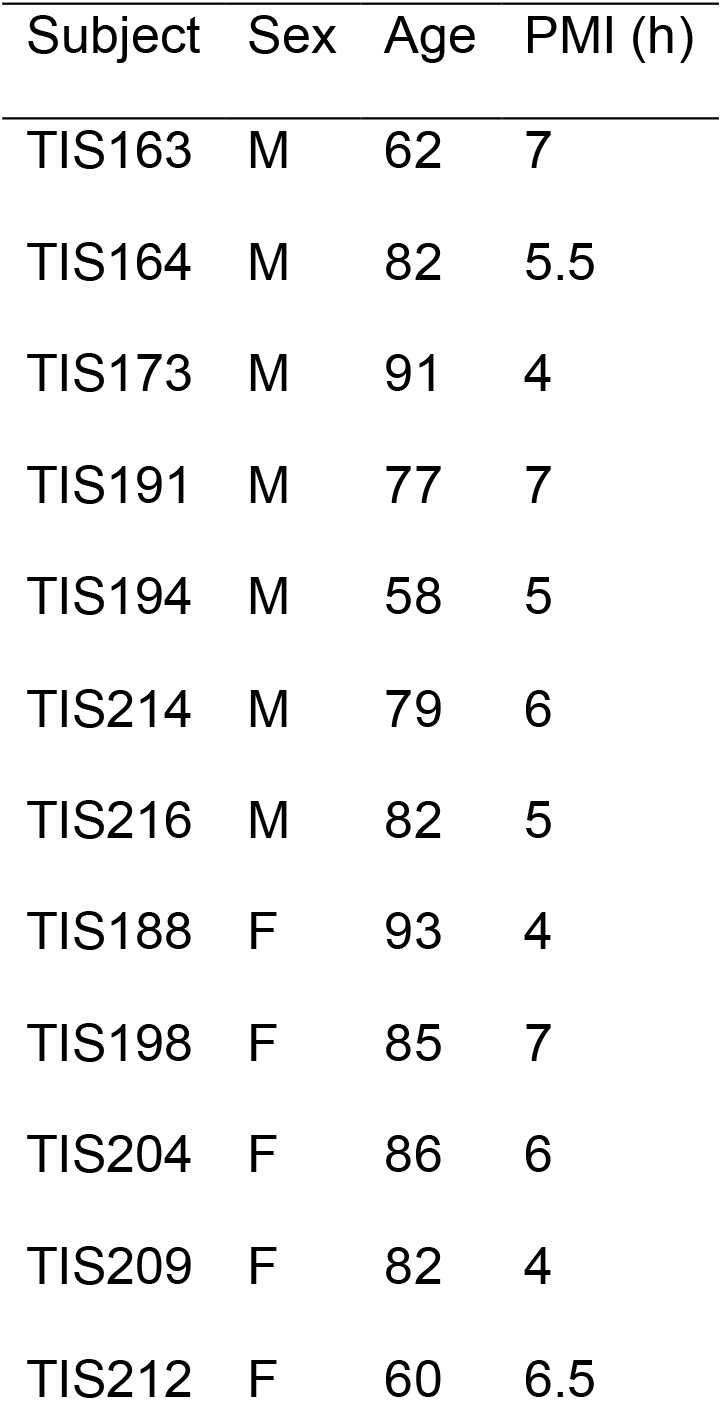
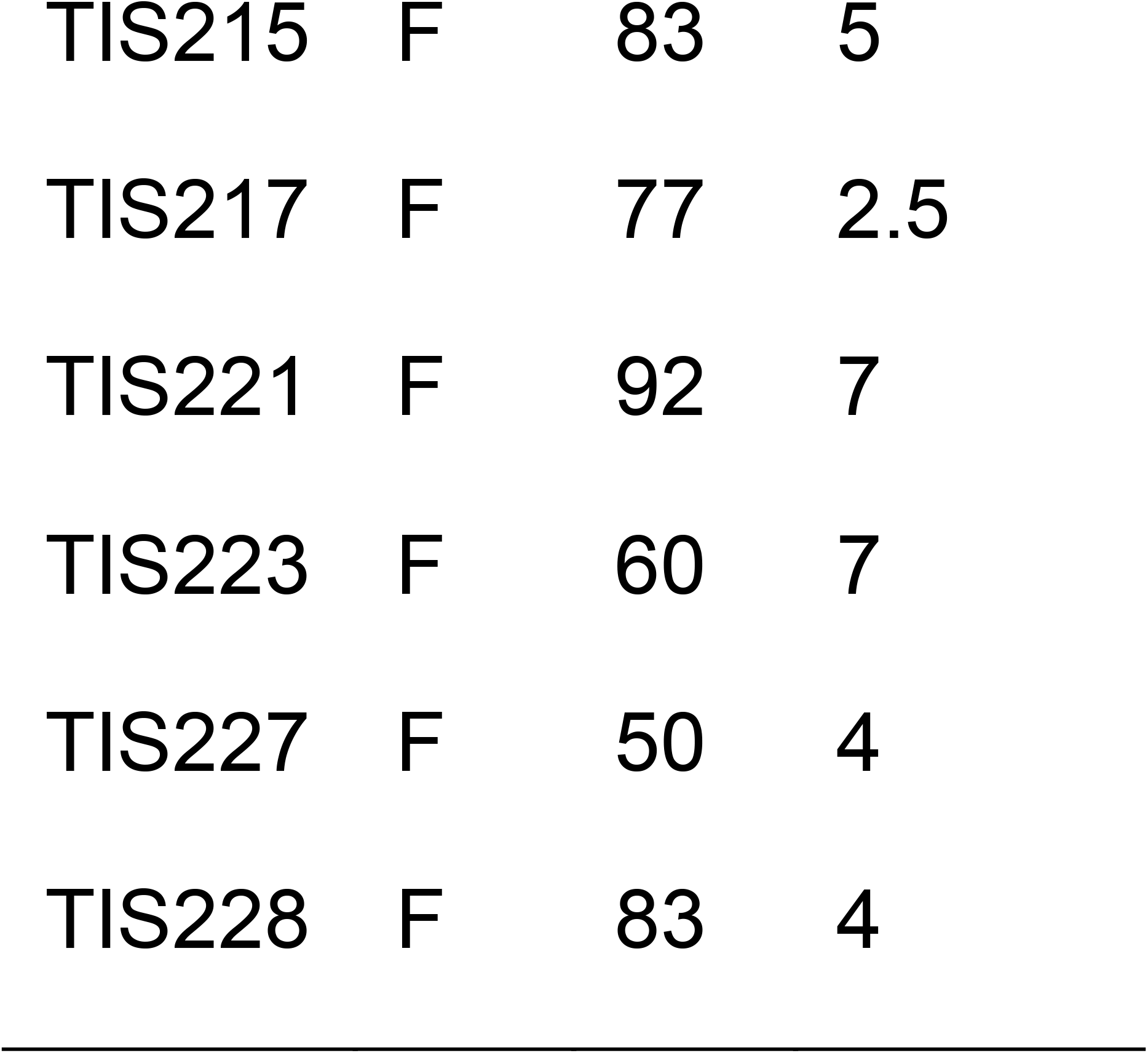
Kinome array subject demographics for the AD cohort (control subjects only).

**Supplementary Table S3.**
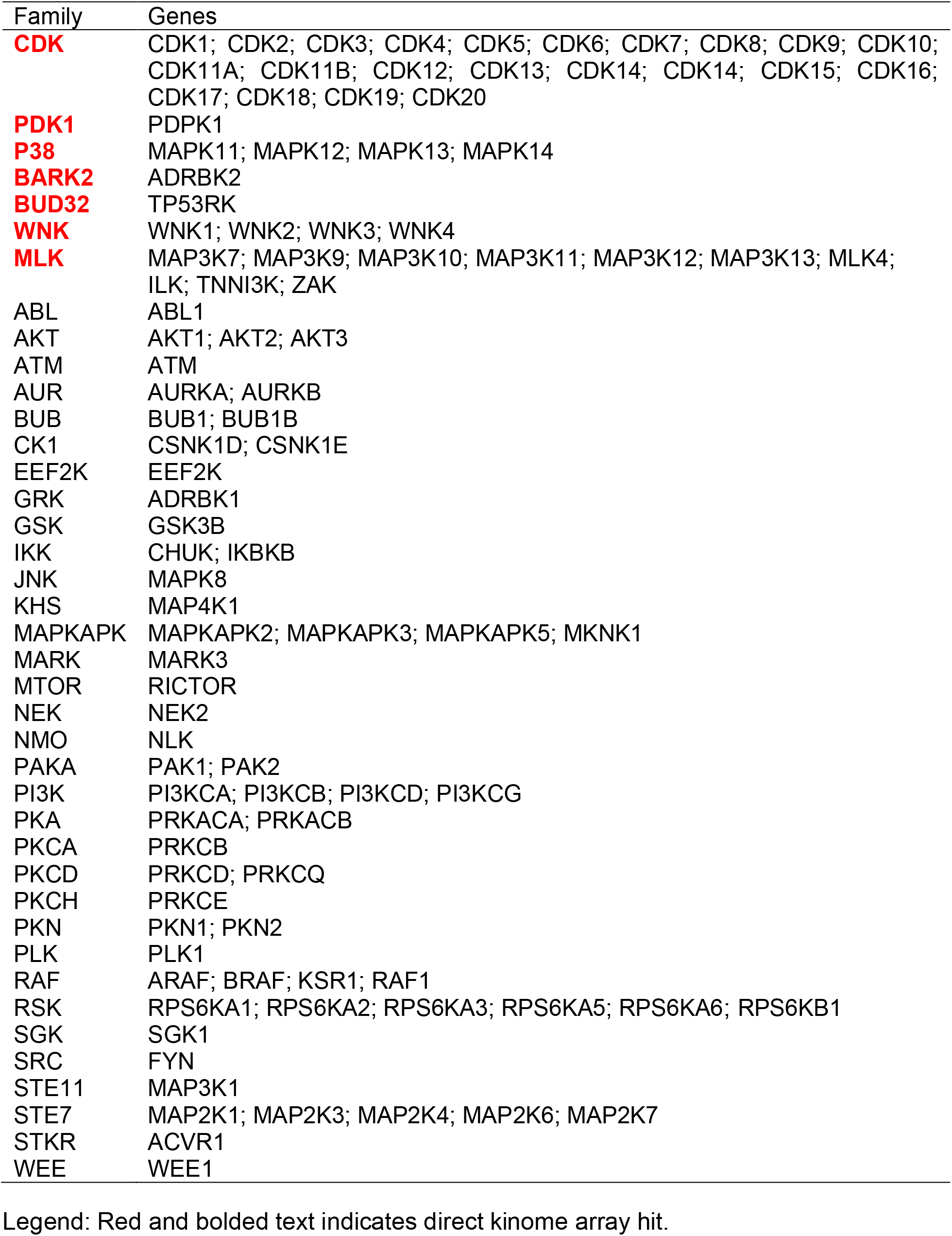
Female vs. male DLPFC kinase network gene list.

| Family | Genes |
| --- | --- |
| **CDK** | CDK1; CDK2; CDK3; CDK4; CDK5; CDK6; CDK7; CDK8; CDK9; CDK10; CDK11A; CDK11B; CDK12; CDK13; CDK14; CDK14; CDK15; CDK16; CDK17; CDK18; CDK19; CDK20 |
| **PDK1** | PDPK1 |
| **P38** | MAPK11; MAPK12; MAPK13; MAPK14 |
| **BARK2** | ADRBK2 |
| **BUD32** | TP53RK |
| **WNK** | WNK1; WNK2; WNK3; WNK4 |
| **MLK** | MAP3K7; MAP3K9; MAP3K10; MAP3K11; MAP3K12; MAP3K13; MLK4; ILK; TNNI3K; ZAK |
| ABL | ABL1 |
| AKT | AKT1; AKT2; AKT3 |
| ATM | ATM |
| AUR | AURKA; AURKB |
| BUB | BUB1; BUB1B |
| CK1 | CSNK1D; CSNK1E |
| EEF2K | EEF2K |
| GRK | ADRBK1 |
| GSK | GSK3B |
| IKK | CHUK; IKBKB |
| JNK | MAPK8 |
| KHS | MAP4K1 |
| MAPKAPK | MAPKAPK2; MAPKAPK3; MAPKAPK5; MKNK1 |
| MARK | MARK3 |
| MTOR | RICTOR |
| NEK | NEK2 |
| NMO | NLK |
| PAKA | PAK1; PAK2 |
| PI3K | PI3KCA; PI3KCB; PI3KCD; PI3KCG |
| PKA | PRKACA; PRKACB |
| PKCA | PRKCB |
| PKCD | PRKCD; PRKCQ |
| PKCH | PRKCE |
| PKN | PKN1; PKN2 |
| PLK | PLK1 |
| RAF | ARAF; BRAF; KSR1; RAF1 |
| RSK | RPS6KA1; RPS6KA2; RPS6KA3; RPS6KA5; RPS6KA6; RPS6KB1 |
| SGK | SGK1 |
| SRC | FYN |
| STE11 | MAP3K1 |
| STE7 | MAP2K1; MAP2K3; MAP2K4; MAP2K6; MAP2K7 |
| STKR | ACVR1 |
| WEE | WEE1 |

Legend: Red and bolded text indicates direct kinome array hit.

**Supplementary Table S4.** Gene expression changes in kinases emerging as direct kinome array hits in cell-level and region-level transcriptome databases of female vs. male frontal cortex.

|  |  | CELL-LEVEL |  |  |  | REGION-LEVEL |  |  |  |
| --- | --- | --- | --- | --- | --- | --- | --- | --- | --- |
|  |  | Female vs. Male DLPFC LCM Superficial Neurons Microarray |  | Female vs. Male DLPFC LCM Deep Neurons Microarray |  | Female vs. Male Prefrontal cortex Microarray (Xu et al., 2014) |  | Female vs. Male Frontal cortex Microarray (Trabzuni et al., 2013) |  |
| Family | Gene | Log2 FC | P-value | Log2 FC | P-value | Log2 FC | P-value | Log2 FC | Probes |
| CDK | CDK1 | -0.087 | 0.697 | **-0.395** | **0.348** | - | - | 0.000 | 2 |
|  | CDK2 | -0.287 | 0.290 | **0.388** | **0.119** | - | - | 0.013 | 2 |
|  | CDK3 | -0.048 | 0.731 | 0.169 | 0.337 | 0.011 | 0.876 | 0.029 | 1 |
|  | CDK4 | 0.051 | 0.714 | 0.065 | 0.713 | 0.111 | 0.386 | 0.030 | 1 |
|  | CDK5 | **-0.360** | **0.067** | -0.280 | 0.416 | **-0.342** | **0.197** | -0.014 | 1 |
|  | CDK6 | 0.287 | 0.050 | 0.215 | 0.265 | 0.219 | 0.335 | -0.028 | 1 |
|  | CDK7 | **-0.938** | **0.053** | **0.571** | **0.276** | -0.156 | 0.267 | -0.056 | 1 |
|  | CDK8 | **0.338** | **0.353** | -0.170 | 0.501 | -0.024 | 0.698 | -0.042 | 1 |
|  | CDK9 | 0.071 | 0.708 | 0.011 | 0.959 | 0.040 | 0.695 | 0.122 | 1 |
|  | CDK10 | 0.129 | 0.688 | **-0.349** | **0.247** | 0.138 | 0.372 | 0.055 | 2 |
|  | CDK11A | 0.075 | 0.777 | -0.154 | 0.679 | - | - | 0.107 | 1 |
|  | CDK11B | 0.001 | 0.999 | **-1.828** | **0.132** | - | - | 0.005 | 1 |
|  | CDK12 | -0.228 | 0.097 | -0.072 | 0.694 | - | - | -0.035 | 1 |
|  | CDK13 | -0.026 | 0.915 | **0.455** | **0.050** | - | - | 0.054 | 2 |
|  | CDK14 | -0.187 | 0.282 | -0.270 | 0.383 | - | - | -0.116 | 1 |
|  | CDK15 | -0.283 | 0.222 | -0.271 | 0.199 | - | - | -0.005 | 2 |
|  | CDK16 | -0.245 | 0.394 | **-0.797** | **0.305** | - | - | -0.028 | 2 |
|  | CDK17 | -0.284 | 0.177 | **-0.774** | **0.017** | - | - | 0.030 | 1 |
|  | CDK18 | -0.279 | 0.535 | 0.205 | 0.757 | - | - | 0.001 | 4 |
|  | CDK19 | 0.177 | 0.643 | -0.079 | 0.864 | - | - | 0.009 | 1 |
|  | CDK20 | -0.212 | 0.325 | -0.029 | 0.898 | - | - | 0.013 | 2 |
| PDK1 | PDPK1 | 0.242 | 0.220 | -0.020 | 0.934 | -0.005 | 0.971 | -0.008 | 5 |
| P38 | MAPK11 | 0.119 | 0.500 | 0.292 | 0.347 | -0.031 | 0.838 | -0.016 | 2 |
|  | MAPK12 | -0.123 | 0.396 | **0.389** | **0.032** | -0.003 | 0.982 | 0.020 | 1 |
|  | MAPK13 | -0.023 | 0.935 | -0.095 | 0.664 | **-0.330** | **0.195** | -0.013 | 1 |
|  | MAPK14 | **0.506** | **0.083** | 0.001 | 0.997 | - | - | -0.033 | 4 |
| BARK2 | ADRBK2 | 0.001 | 0.996 | -0.213 | 0.474 | - | - | - | - |
| BUD32 | TP53RK | 0.043 | 0.880 | **-0.317** | **0.135** | -0.068 | 0.399 | -0.051 | 1 |
| WNK | WNK1 | -0.057 | 0.919 | 0.199 | 0.685 | **0.366** | **0.255** | 0.012 | 3 |
|  | WNK2 | **0.357** | **0.379** | **0.463** | **0.016** | -0.005 | 0.970 | 0.079 | 1 |
|  | WNK3 | -0.242 | 0.215 | -0.103 | 0.529 | -0.002 | 0.975 | -0.005 | 3 |
|  | WNK4 | 0.223 | 0.359 | -0.071 | 0.782 | - | - | -0.008 | 1 |
| MLK | MAP3K7 | 0.153 | 0.395 | -0.023 | 0.942 | -0.187 | 0.198 | -0.012 | 2 |
|  | MAP3K9 | 0.149 | 0.433 | **-0.353** | **0.158** | **-0.357** | **0.242** | 0.004 | 1 |
|  | MAP3K10 | 0.157 | 0.575 | 0.067 | 0.730 | 0.124 | 0.570 | -0.019 | 1 |
|  | MAP3K11 | **0.304** | **0.281** | -0.206 | 0.605 | 0.107 | 0.672 | 0.070 | 1 |
|  | MAP3K12 | **-0.339** | **0.142** | **0.395** | **0.353** | -0.012 | 0.936 | 0.095 | 1 |
|  | MAP3K13 | -0.235 | 0.498 | 0.029 | 0.926 | 0.115 | 0.447 | 0.000 | 1 |
|  | MLK4 | 0.110 | 0.550 | 0.151 | 0.512 | - | - | - | - |
|  | ILK | 0.078 | 0.764 | -0.003 | 0.994 | 0.203 | 0.216 | 0.000 | 3 |
|  | TNNI3K | 0.052 | 0.761 | 0.030 | 0.899 | -0.236 | 0.245 | 0.038 | 1 |
|  | ZAK | -0.210 | 0.271 | 0.073 | 0.795 | **0.304** | **0.188** | 0.035 | 3 |

Legend: Red and bolded text indicates genes with log2 FC > 0.3, log2 FC < -0.3, or significant findings (p < 0.05) irrespective of FC values.

**Supplementary Table S5.** Gene expression changes in kinases emerging as indirect kinome array hits (i.e. connected via STRING with the direct kinome array hits) in cell-level and region-level transcriptome databases of female vs. male frontal cortex.

|  |  | CELL-LEVEL |  |  |  | REGION-LEVEL |  |  |  |
| --- | --- | --- | --- | --- | --- | --- | --- | --- | --- |
|  |  | Female vs. Male DLPFC LCM Superficial Neurons Microarray |  | Female vs. Male DLPFC LCM Deep Neurons Microarray |  | Female vs. Male Prefrontal cortex Microarray (Xu et al., 2014) |  | Female vs. Male Frontal cortex Microarray (Trabzuni et al., 2013) |  |
| Family | Gene | Log2 FC | P-value | Log2 FC | P-value | Log2 FC | P-value | Log2 FC | Probes |
| ABL | ABL1 | -0.270 | 0.298 | 0.247 | 0.457 | 0.158 | 0.247 | 0.015 | 3 |
| AKT | AKT1 | 0.027 | 0.921 | **-0.608** | **0.062** | 0.243 | 0.249 | 0.060 | 3 |
|  | AKT2 | -0.236 | 0.465 | **0.356** | **0.508** | - | - | -0.012 | 1 |
|  | AKT3 | **-1.221** | **0.188** | **-0.513** | **0.570** | 0.023 | 0.526 | 0.008 | 4 |
| ATM | ATM | -0.101 | 0.753 | **-0.472** | **0.219** | - | - | -0.026 | 3 |
| AUR | AURKA | **0.438** | **0.233** | **-0.786** | **0.054** | -0.051 | 0.349 | -0.004 | 2 |
|  | AURKB | **-0.626** | **0.025** | 0.091 | 0.776 | - | - | 0.007 | 1 |
| BUB | BUB1 | -0.134 | 0.687 | 0.115 | 0.681 | - | - | 0.002 | 1 |
|  | BUB1B | -0.285 | 0.264 | -0.130 | 0.517 | 0.005 | 0.927 | 0.020 | 1 |
| CK1 | CSNK1D | 0.049 | 0.732 | **0.308** | **0.087** | -0.071 | 0.318 | -0.041 | 2 |
|  | CSNK1E | 0.004 | 0.990 | -0.052 | 0.858 | 0.131 | 0.303 | 0.008 | 3 |
| EEF2K | EEF2K | 0.029 | 0.864 | 0.131 | 0.458 | 0.177 | 0.374 | 0.012 | 1 |
| GRK | ADRBK1 | -0.105 | 0.706 | 0.281 | 0.395 | - | - | - | - |
| GSK | GSK3B | -0.168 | 0.414 | **-0.767** | **0.005** | 0.054 | 0.748 | 0.099 | 1 |
| IKK | CHUK | **-0.428** | **0.257** | 0.146 | 0.762 | 0.060 | 0.529 | 0.001 | 1 |
|  | IKBKB | 0.117 | 0.465 | -0.063 | 0.709 | 0.118 | 0.203 | 0.046 | 2 |
| JNK | MAPK8 | **-0.305** | **0.379** | **-0.394** | **0.279** | -0.005 | 0.917 | 0.008 | 2 |
| KHS | MAP4K1 | 0.035 | 0.867 | -0.057 | 0.873 | 0.010 | 0.807 | 0.058 | 2 |
| MAPKAPK | MAPKAPK2 | -0.084 | 0.562 | 0.270 | 0.128 | - | - | 0.018 | 2 |
|  | MAPKAPK3 | 0.204 | 0.290 | 0.174 | 0.381 | **0.459** | **0.234** | 0.027 | 1 |
|  | MAPKAPK5 | 0.203 | 0.207 | -0.084 | 0.721 | -0.039 | 0.616 | -0.063 | 2 |
|  | MKNK1 | 0.117 | 0.447 | 0.116 | 0.485 | - | - | 0.055 | 2 |
| MARK | MARK3 | -0.041 | 0.872 | 0.137 | 0.714 | -0.113 | 0.209 | 0.033 | 1 |
| MTOR | RICTOR | -0.117 | 0.506 | -0.223 | 0.288 | - | - | 0.027 | 1 |
| NEK | NEK2 | **-0.377** | **0.107** | **0.319** | **0.290** | -0.035 | 0.693 | 0.020 | 2 |
| NMO | NLK | **-0.770** | **0.008** | **-0.613** | **0.069** | -0.298 | 0.243 | -0.079 | 1 |
| PAKA | PAK1 | **-0.492** | **0.059** | **-0.992** | **0.043** | **-0.670** | **0.188** | -0.038 | 1 |
|  | PAK2 | **0.381** | **0.139** | 0.001 | 0.996 | 0.083 | 0.368 | 0.015 | 2 |
| PI3K | PI3KCA | **0.428** | **0.221** | **-0.419** | **0.182** | - | - | -0.003 | 2 |
|  | PI3KCB | -0.123 | 0.586 | 0.049 | 0.876 | -0.194 | 0.365 | -0.091 | 1 |
|  | PI3KCD | 0.061 | 0.834 | 0.233 | 0.518 | -0.034 | 0.790 | 0.056 | 1 |
|  | PI3KCG | -0.174 | 0.314 | -0.162 | 0.498 | - | - | 0.034 | 2 |
| PKA | PRKACA | **0.646** | **0.226** | 0.179 | 0.738 | 0.001 | 0.991 | -0.001 | 2 |
|  | PRKACB | **-0.351** | **0.113** | **-0.584** | **0.317** | - | - | -0.041 | 5 |
| PKCA | PRKCB | **-0.407** | **0.155** | **-1.263** | **0.003** | **-0.503** | **0.250** | -0.086 | 3 |
| PKCD | PRKCD | **0.722** | **0.058** | **0.566** | **0.016** | - | - | -0.009 | 2 |
|  | PRKCQ | 0.213 | 0.195 | -0.196 | 0.291 | -0.227 | 0.316 | -0.106 | 1 |
| PKCH | PRKCE | **-0.596** | **0.011** | **-0.401** | **0.198** | **-0.551** | **0.197** | -0.042 | 1 |
| PKN | PKN1 | **-0.304** | **0.353** | **-0.618** | **0.060** | - | - | 0.026 | 4 |
|  | PKN2 | **0.361** | **0.190** | 0.160 | 0.477 | 0.071 | 0.345 | 0.091 | 1 |
| PLK | PLK1 | **0.589** | **0.291** | 0.070 | 0.929 | - | - | 0.010 | 1 |
| RAF | ARAF | -0.244 | 0.233 | -0.079 | 0.840 | 0.255 | 0.207 | 0.043 | 1 |
|  | BRAF | 0.018 | 0.904 | -0.296 | 0.136 | - | - | -0.042 | 2 |
|  | KSR1 | **0.359** | **0.243** | -0.221 | 0.468 | - | - | 0.010 | 1 |
|  | RAF1 | 0.247 | 0.203 | 0.065 | 0.792 | 0.042 | 0.570 | 0.072 | 1 |
| RSK | RPS6KA1 | **-0.517** | **0.511** | **1.728** | **0.012** | -0.035 | 0.756 | -0.002 | 2 |
|  | RPS6KA2 | **0.413** | **0.203** | **0.561** | **0.222** | 0.153 | 0.203 | 0.015 | 3 |
|  | RPS6KA3 | **1.152** | **0.037** | **-0.382** | **0.245** | 0.024 | 0.573 | -0.002 | 4 |
|  | RPS6KA5 | 0.237 | 0.411 | -0.132 | 0.632 | - | - | -0.042 | 2 |
|  | RPS6KA6 | 0.218 | 0.109 | -0.145 | 0.400 | - | - | 0.002 | 4 |
|  | RPS6KB1 | 0.273 | 0.580 | **-1.161** | **0.009** | -0.025 | 0.811 | -0.006 | 1 |
| SGK | SGK1 | **-0.526** | **0.471** | **-1.082** | **0.094** | -0.088 | 0.820 | -0.014 | 2 |
| SRC | FYN | -0.168 | 0.626 | -0.036 | 0.944 | 0.142 | 0.270 | 0.022 | 4 |
| STE11 | MAP3K1 | 0.193 | 0.228 | 0.091 | 0.738 | **0.385** | **0.189** | 0.054 | 1 |
| STE7 | MAP2K1 | -0.047 | 0.831 | **-0.310** | **0.201** | **-0.585** | **0.203** | -0.142 | 1 |
|  | MAP2K3 | 0.148 | 0.399 | **0.625** | **0.015** | - | - | 0.002 | 6 |
|  | MAP2K4 | **0.497** | **0.133** | 0.079 | 0.845 | **-0.406** | **0.189** | -0.076 | 2 |
|  | MAP2K6 | **0.323** | **0.287** | -0.007 | 0.985 | - | - | -0.034 | 1 |
|  | MAP2K7 | **0.368** | **0.086** | 0.088 | 0.642 | 0.053 | 0.560 | 0.042 | 1 |
| STKR | ACVR1 | **-0.722** | **0.028** | -0.092 | 0.786 | **0.393** | **0.188** | -0.042 | 1 |
| WEE | WEE1 | 0.225 | 0.455 | 0.162 | 0.371 | - | - | 0.024 | 1 |

Legend: Red and bolded text indicates genes with $\log_2 \text{FC} > 0.3$ , $\log_2 \text{FC} < -0.3$ , or significant findings ( $p < 0.05$ ) irrespective of FC values.

